# Comparison of real-time quantitative Polymerase Chain Reaction (PCR) and digital droplet PCR for quantification of hormone membrane receptor *FSHR, GPER* and *LHCGR* transcripts in human primary granulosa lutein cells

**DOI:** 10.1101/2020.06.22.164434

**Authors:** Samantha Sperduti, Claudia Anzivino, Maria Teresa Villani, Gaetano De Feo, Manuela Simoni, Livio Casarini

## Abstract

**Background:** Quantitative real time polymerase chain reaction (qPCR) and droplet digital PCR (ddPCR) are methods used for gene expression analysis in several contexts, including reproductive endocrinology.

**Objectives:** Herein, we compared qPCR and ddPCR technologies for gene expression analysis of hormone membrane receptor-encoding genes, such as follicle-stimulating hormone (*FSHR)*, G protein-coupled estrogen (*GPER)* and choriogonadotropin receptors (LHCGR), as well as the commonly used *RPS7* housekeeping gene, in order to identify the most reliable method to be applied for gene expression analysis in the context of human reproduction.

**Methods:** Total RNA was extracted from human primary granulosa cells of donor patients undergoing assisted reproduction and used for gene expression analysis by qPCR and ddPCR, after finding the optimal annealing temperature.

**Results:** Both techniques provided results reflecting the low number of *FSHR* and *GPER* transcripts, although ddPCR detected also unspecific transcripts in using *RPS7* primers and quantified the low-expressed genes with major accuracy, thanks to its higher reaction efficiency. The absolute *FSHR* and *GPER* transcript number was also determined by ddPCR, resulting in 50- to 170-fold lower amount than *LHCGR* transcripts.

**Conclusion:** These results suggest that ddPCR is the candidate technology for analysis of genes with relatively low expression levels and provides useful insights for characterizing hormone receptor expression levels in the context of reproductive endocrinology.

## Introduction

Gene expression analysis may be applied to the context of reproductive endocrinology, where it helps to understand hormone-specific functions in target cells of hormones regulating human reproduction. Ovarian function is regulated by two gonadotropins, the luteinizing hormone (LH) and the follicle-stimulating hormone (FSH), as well as sex steroid hormones such as estrogens. These molecules act through specific receptors expressed on gonadal target cells, i.e. FSH (FSHR) (Simoni et al., 1997), LH and choriogonadotropin (LHCGR) (Casarini et al., 2018) and G protein-coupled estrogen receptor (GPER) (Revankar et al., 2005), which induce the activation of different intracellular signalling pathways, depending on their expression levels on the cell surface (Casarini et al., 2016a; Zhu et al., 1994). Therefore, the determination of individual-specific *FSHR, LHCGR* and *GPER* expression levels is relevant for the study of human reproduction pathophysiology and drug effects (Regard et al., 2008).

Quantitative real time polymerase chain reaction (qPCR) represented for a long time the gold standard method for assessing gene expression, thanks to the use of fluorophores (Kubista et al., 2006; Morrison et al., 1998). The relative quantification of the target is determined by monitoring its amplification in real time during the PCR, after plotting the florescence against the number of PCR cycles on a logarithmic scale; the lowest cycle number at which the fluorescence emitted by DNA amplification is higher than that of the fixed threshold for background signal is defined as “quantification cycle” (Cq) (Kubista et al., 2006). The amount of DNA copies should ideally be double at each cycle, allowing the determination of the reaction efficiency by creating a standard curve of the Cq changes. This experiment may be executed via titration by serial dilution of the DNA template and plotting the data using linear regression. Efficiency is indicated by the slope, while the quantification of the transcript may be obtained by comparing the Cq with that of a housekeeping gene (Livak and Schmittgen, 2001). To reach reproducible and reliable results it is necessary to optimize each PCR reaction. Inaccurate primer design and poorly optimized qPCR experiments may, in fact, influence data interpretation, leading to misunderstanding of their biological significance (Bustin et al., 2009; Tellinghuisen and Spiess, 2015).

Droplet digital PCR (ddPCR) is a recently developed technology, providing several advantages in comparison to qPCR. Briefly, ddPCR is a fluorescence based assay, allowing partitioning of each single gene copy into one of the thousands nanoliter-sized droplets, which are subjected to end-point PCR amplification (Beer et al., 2008). Thus, each reaction is analysed to determine the number of PCR-positive droplets, containing the amplified target, over the negative droplets, and the number of target copies in the original sample is inferred by Poisson statistical analysis (Hindson et al., 2013). Moreover, ddPCR allows a robust and powerful end-point quantification of targets without the use of a standard curve (Bizouarn, 2014; Hindson et al., 2011; Sanders et al., 2011). This technology is characterized by high sensitivity, accuracy, technical precision, reproducibility and improved tolerance to PCR inhibitors, providing many advantages compared to qPCR, especially for low target quantifications (Dingle et al., 2013).

In this study, we validated *FSHR, GPER* and *LHCGR* gene expression by comparing qPCR and ddPCR results, in order to identify the most reliable method to be applied for gene expression analysis in the context of human reproduction. The expression of the housekeeping *RPS7* gene (Casarini et al., 2017, 2016b, 2016a) was also evaluated.

## Materials and Methods

### Granulosa cells isolation and ribonucleic acid (RNA) extraction

Sample RNAs were extracted from human primary granulosa lutein cells (hGLC). They were recovered from follicular fluid aspirate from six women undergoing oocyte retrieval for assisted reproduction, as previously described (Casarini et al., 2014, 2012), and pooled in two groups. These patients were diagnosed as infertile due to tubal or male factor and were 25-45 years old, without endocrine abnormalities and severe viral or bacterial infections. Written consent was collected from women under local Ethics Committee permission (Nr. 796 19th June 2014, Reggio Emilia, Italy). hGLC were grown at 37°C and 5.0% CO_2_ in DMEM/F12 medium, supplemented with 10.0% FBS, 2.0 mM L-glutamine, 100.0 IU/ml penicillin, 0.1 mg/ml streptomycin (all from Thermo Fisher Scientific, Waltham, MA, USA) and 250.0 ng/ml Fungizone (Merck KGaA, Darmstadt, Germany). After 6 days of culturing, required for recovering the expression of hormone receptors (Nordhoff et al., 2011), total RNA was extracted from pools of 2500000 cells using the EZ1 RNA Tissue Mini Kit (#959034, Qiagen, Hilden, Germany), according to the manufacturer’s instructions, and quantified by a Nanodrop 2000 spectrophotometer (Thermo Fisher Scientific). 17.5 pg/cell RNA were obtained.

### cDNA synthesis and primer design

Reaction mixes for complementary DNA (cDNA) synthesis were assembled in 20.0 µl each, using the high-capacity cDNA reverse transcription kit (#4374966, Applied Biosystems, Foster City, CA, USA), as previously described (Cantanelli et al., 2014). Briefly, 10 µl of 2x reverse transcription mix, containing 2 µl of 10x RT Buffer, 2 µl of 10x RT Random Primers, 0.8 µl of 25x dNTP Mix (100.0 mM), 1 µl of 20.0 U/µl RNase inhibitor, 1 µl of 50.0 U/µl MultiScribe™ Reverse Transcriptase and 3.2 µl of nuclease-free water, were prepared on ice. 10.0 µl of the reverse transcription master mix was added to reaction tubes containing 2.0 µg of RNA previously diluted in 10.0 µl of RNAse-free water each. Reverse transcription reactions were performed using a C1000 Thermal cycler (Bio-Rad Laboratories Inc., Hercules, CA, USA) under the following conditions: 25.0°C for 10 minutes and 37.0°C for 120 minutes. The expression of *FSHR, GPER* and *LHCGR* genes were assessed using specific primer probes and the *RPS7* housekeeping gene expression was used as a reference (Casarini et al., 2017, 2016a, 2016b). Primers used for both qPCR and ddPCR (table 1) were designed using the Primer3 web-based tool (http://www.ncbi.nlm.nih.gov/tools/primer-blast/).

**Table 1.**
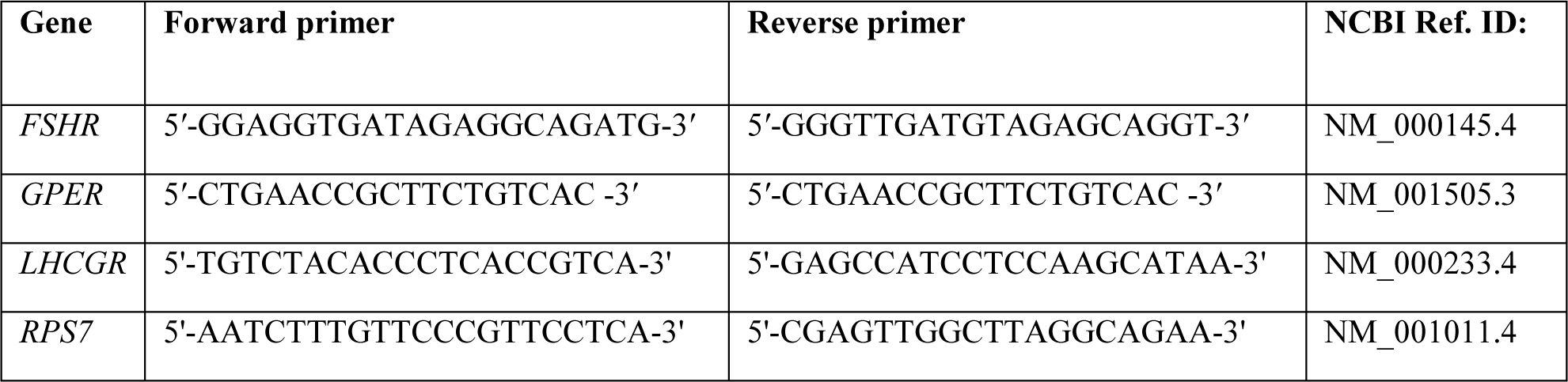
Primer sequences used for *FSHR, GPER, LHCGR* and *RPS7* gene expression analysis.

### qPCR and ddPCR reactions setup

qPCR and ddPCR reactions were performed in duplicate, using the CFX96™ Real-Time PCR Detection System and the QX200 Droplet Digital PCR (ddPCR™) System (Bio-Rad Laboratories Inc.), respectively. For each primer pair, 20 µl of a unique reaction mixture containing nuclease-free water, forward and reverse primer probes (at final concentration 125 nM each) and cDNA were prepared. Specifically, 5.0 ng of cDNA were used to evaluate the most suitable annealing temperature, while serial dilutions of the stock cDNA (0.5-100.0 ng/µl range) were performed for assessing primer efficiency and sensibility. The mixture was divided into two reaction tubes and 10 µl of the specific SYBR dye-based master mix, SsoFast™ EvaGreen® Supermix for qPCR (#1725202) and QX200™ ddPCR™ EvaGreen Supermix (#186-4034) for ddPCR (both from Bio-Rad Laboratories Inc.), was added. ddPCR droplets were generated on the QX200™ Droplet Generator (Bio-Rad Laboratories Inc.) and amplification reactions were performed according to these settings: 5 min enzyme activation at 95°C; 40 cycles of 30 s each for DNA denaturation at 95°C, 60 s for primer annealing and extension at 56.0, 58.0 or 60.0°C, as indicated in the results section. Signals were stabilized 5 min at 4°C followed by 5 min at 90°C. All temperature variations were performed with a ramp rate of 2°C/s. Positive droplets were determined using the QX200™ Droplet Reader (Bio-Rad Laboratories Inc.). For qPCR analysis, reaction conditions were as follows: 30 s enzyme activation at 95°C; 45 cycles of 3 s each for DNA denaturation at 95°C, 5 s primer annealing and extension at 56.0, 58.0 or 60.0°C. After qPCR cycling, a melt curve from 60.0°C to 95.0°C with increments of 0.5°C and 5 s hold was applied.

### Agarose gel electrophoresis

Primer specificity was validated running the PCR products obtained by qPCR on a 3.0% agarose-gel electrophoresis, assessing amplicons lengths. Images were acquired by the QuantityOne analysis software (Bio-Rad Laboratories Inc., Hercules, CA, USA).

### Data processing and statistical analysis

Relative gene expression values (Cq) were obtained by qPCR and processed using the CFX Manager (v.3.1) software (Bio-Rad Laboratories Inc.). ddPCR data were analysed by QuantaSoft (v.1.7.4) software (Bio-Rad Laboratories Inc.) and only reactions with a number of total droplets >13000 per 20 µl were considered. The absolute quantification of cDNA was expressed as target transcripts per 1.0 µl. Relative fold differences in gene expression levels were expressed as 2^-ΔCq^, where ΔCq value resulted by data normalization over Cq values obtained using the maximum concentration of template cDNA. The coefficient of variation % (%CV) has been used to assess the variability, as previously described (Taylor et al., 2017). Primer efficiency percentage was calculated, after interpolation by linear regression of the target gene amount plotted against the cDNA template amount, as follows: *E*=(10^−1/(Slope value)^-1)x100. Graphical elaborations of efficiency data from ddPCR analysis, as well as r^2^ and slope calculation were performed by GraphPad Prism 6.0 software (GraphPad Software Inc., San Diego, CA, USA).

## Results

### Analysis of melting temperature-dependency of qPCR and ddPCR performance

The comparison between ddPCR and qPCR was performed after adjustment for the best primer annealing temperature between 56.0, 58.0 and 60.0°C. qPCR Cq values and ddPCR target copies per µl were extracted from raw data (supplementary figure S1) and reported in a table (table 2). Both qPCR and ddPCR results converged in demonstrating overall poor expression levels of *FSHR* and *GPER*, indicated by relatively high Cq values and low gene copies per µl and %CV. Moreover, qPCR did not detect *GPER* expression when reactions were performed at melting temperature below 60°C. These data were corroborated by <1.0 transcripts copy/µl by ddPCR. Reactions performed using *LHCGR* and *RPS7* primer probes resulted in Cq values ranging between 28.0±0.06 and 33.7±0.1 by qPCR, as well as in 119.0±1.4 and 377±11.3 transcript copies within the temperature range considered.

**Table 2.** *FSHR, GPER, LHCGR* and *RPS7* gene expression levels detected by qPCR and ddPCR under different annealing temperatures. n=number of experiments with successful target amplification (out of 2); SD=standard deviation; NTC=negative control sample; NA=not amplified; UN=unavailable.

|  | qPCR |  |  |  |  |  | ddPCR |  |  |  |  |
| --- | --- | --- | --- | --- | --- | --- | --- | --- | --- | --- | --- |
| Annealing Temperature (°C) | Cq (means) | n | SD | %C V | NTC |  | gene transcripts/μl (means) | n | SD | %C V | NTC |
| ***FSHR*** |  |  |  |  |  |  |  |  |  |  |  |
| 60.0 | 36.2 | 2 | 0.03 | 0.07 | NA |  | 3.1 | 2 | 0.5 | 16.2 | NA |
| 58.2 | 34.7 | 2 | 0.02 | 0.07 | NA |  | 2.5 | 2 | 0.3 | 11.3 | NA |
| 56.0 | 33.8 | 2 | 0.5 | 1.4 | NA |  | 3.0 | 2 | 0.4 | 14.1 | NA |
| ***GPER*** |  |  |  |  |  |  |  |  |  |  |  |
| 60.0 | 45.7 | 1 | UN | UN | NA |  | 1.3 | 2 | 0.1 | 10.9 | NA |
| 58.2 | NA | 0 | NA | NA | NA |  | 0.8 | 2 | 0.1 | 16.8 | NA |
| 56.0 | NA | 0 | NA | NA | NA |  | 0.9 | 2 | 0.5 | 59.5 | NA |
| ***LHCGR*** |  |  |  |  |  |  |  |  |  |  |  |
| 60.0 | 28.5 | 2 | 0.02 | 0.06 | NA |  | 126.0 | 2 | 1.4 | 1.1 | NA |
| 58.2 | 28.0 | 2 | 0.06 | 0.2 | 34.6 |  | 119.0 | 2 | 1.4 | 1.2 | NA |
| 56.0 | 28.0 | 2 | 0.00 | 0.01 | 39.5 |  | 126.0 | 2 | 5.7 | 4.5 | NA |
| ***RPS7*** |  |  |  |  |  |  |  |  |  |  |  |
| 60.0 | 33.7 | 2 | 0.1 | 0.3 | NA |  | 374.0 | 2 | 11.3 | 3.0 | 0.1 |
| 58.2 | 31.9 | 2 | 0.02 | 0.08 | NA |  | 373.5 | 2 | 2.1 | 0.6 | NA |
| 56.0 | 31.4 | 2 | 0.3 | 0.9 | NA |  | 377.0 | 2 | 11.3 | 3.0 | NA |

The specificity of target amplifications was verified by agarose gel electrophoresis after PCR (figure 1) demonstrating the expected amplicons sequence lengths.

**Fig. 1.**
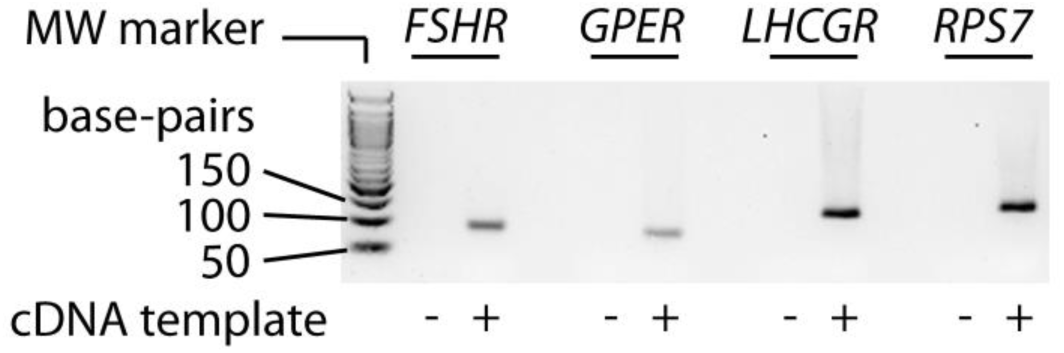
Validation of FSHR, GPER, LHCGR and RPS7 PCR amplicon length. Analysis of PCR fragments by 3% agarose-gel electrophoresis demonstrating amplicon length with expected molecular weights; FSHR=89 bp; GPER=78 bp; LHCGR=118 bp; RPS7=135 bp.

### Evaluation of primer efficiency and sensibility by qPCR and ddPCR

Since the tested melting temperatures produced overall similar results, primer efficiency and sensibility obtained by qPCR and ddPCR were compared using the common melting temperature of 58.0°C. Serial dilutions of the total cDNA (100.0-0.5 ng/µl range) were prepared and the amount of target genes transcripts was measured. Both techniques detected the decrease of transcript copies together with the amount of template cDNA (figure 2, table 3). However, ddPCR data plotting and interpolation by linear regression (not shown) revealed correlation between gene transcripts number (relative fold decrease) and amount of cDNA, resulting in r^2^≥0.99 for all samples, while weaker correlations were found by qPCR data plotting, demonstrated by r^2^ values ranging between 0.86 and 0.95 (linear regression; n=2; p<0.0001).

**Table 3.**
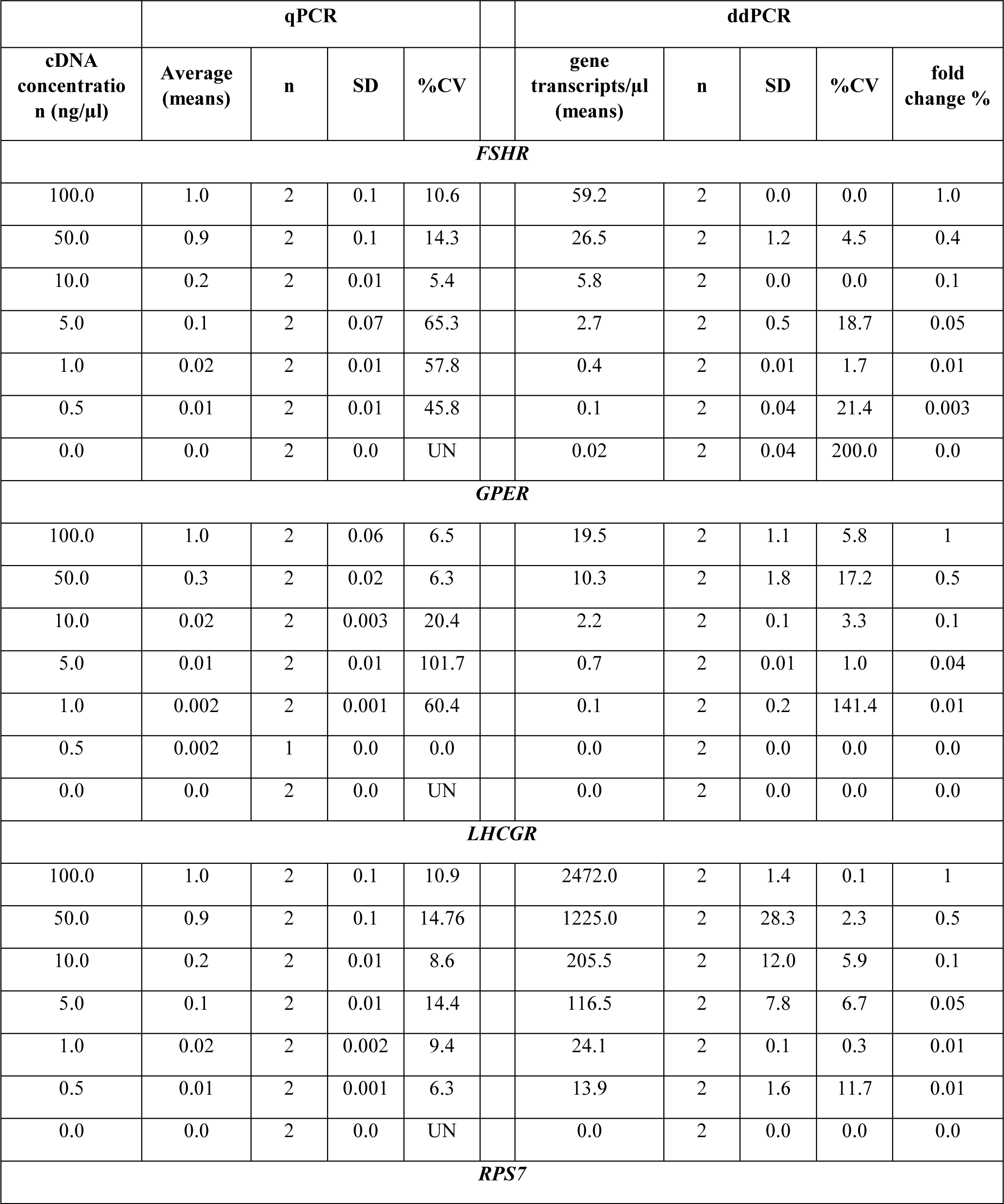
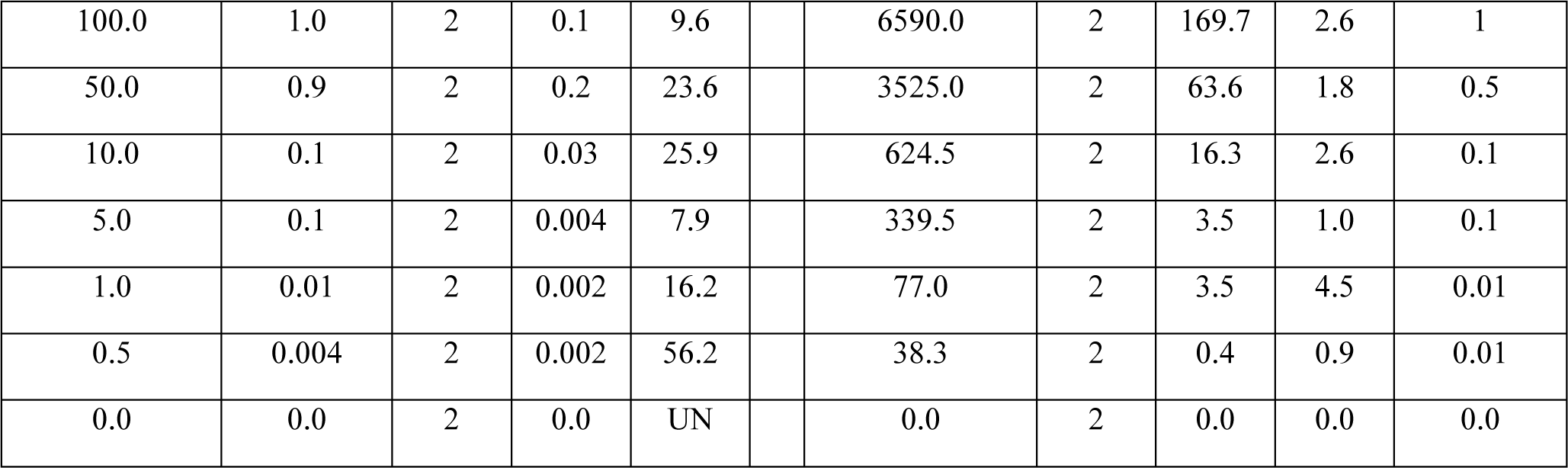
Comparison between qPCR and ddPCR target gene expression analysis. qPCR results (average) are indicated as fold differences of Cqs *versus* the Cq of the sample with maximal template amount (100 ng/ml). ddPCR data are indicated as DNA copy/µl. n=number of experiments with successful target amplification (out of 2); SD=standard deviation; NTC=negative control sample; UN=unavailable.

**Fig. 2.**
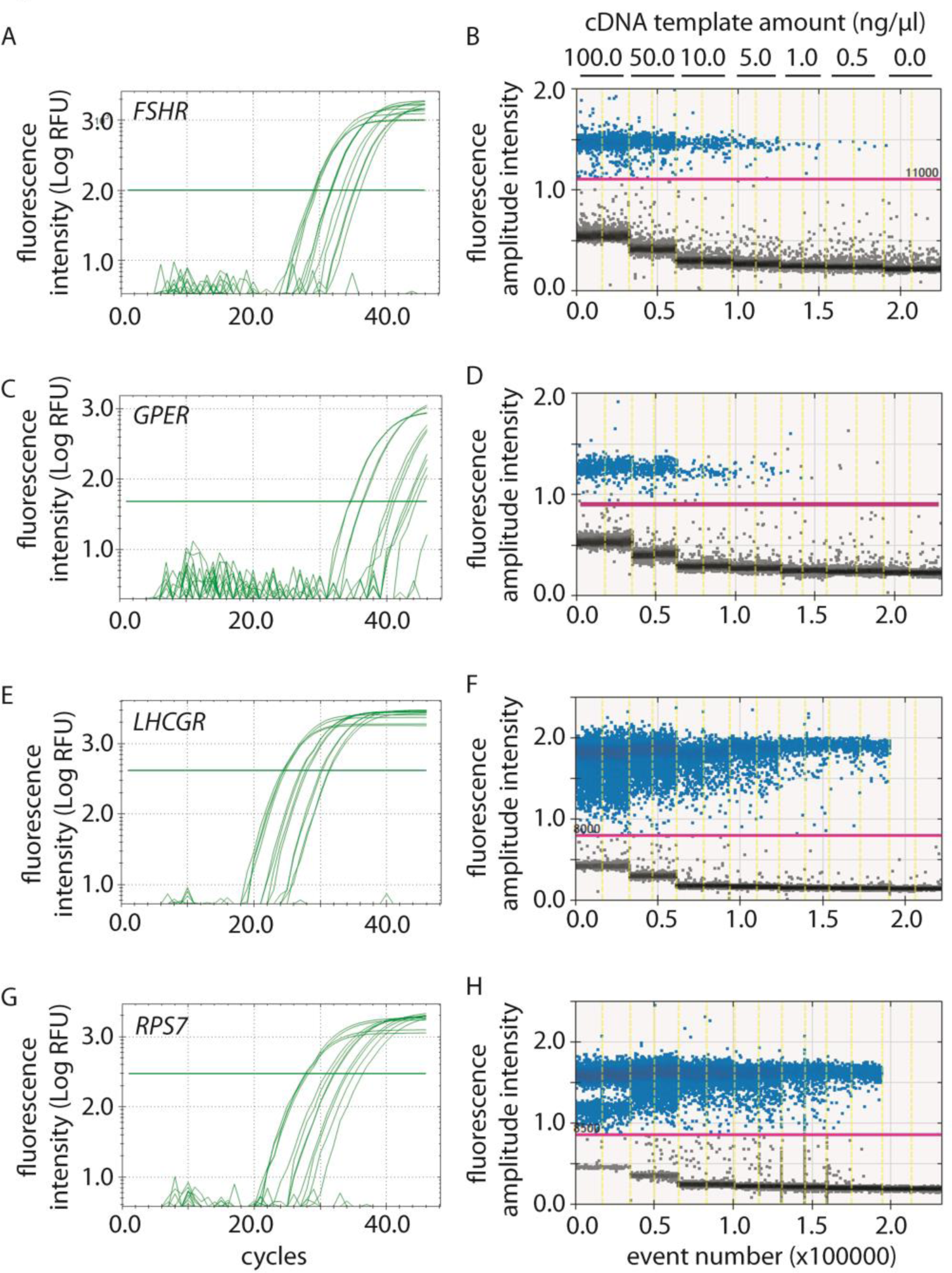
Results of FSHR, GPER, LHCGR and RPS7 transcript amplifications by qPCR and ddPCR. Reactions were performed using scaled cDNA concentrations (100.0 - 0.5 ng/µl cDNA range) using qPCR and ddPCR technologies. A, B) Data from FSHR gene expression analysis. C, D) GPER. E, F) LHCGR. G, H) RPS7. qPCR amplification curves are shown in graphs (left panels) and placed beside ddPCR amplitude plots (right panels).

Variability among ddPCR replicates depends on gene expression levels and resulted in <12% CV for *LHCGR* and *RPS7*, while it was <22% CV for *FSHR* and *GPER* genes (table 3). Different performances were obtained by qPCR, where the variability of replicates was <26% CV at the three highest cDNA concentrations (table 3).

Interestingly, ddPCR analysis using *RPS7* primer probes and 100 ng/µl of template cDNA identified a population of positive droplets with fluorescence amplitude intensity of about 12000 relative light units (RLU), while the *RPS7*-specific signal was at about 16000 RLU (figure 2). Unspecific amplicons were not detected by qPCR.

We considered qPCR to show a better efficiency (*E*) within the range of 80-120%, calculated using linear regression interpolating gene expression data from serial cDNA dilutions in the presence of r^2^≥0.95 (Shehata et al., 2019; Veselenak et al., 2015). While qPCR analysis revealed that *FSHR* and *LHCGR* amplifications were efficient (*E*_*FSHR*_=119.5%; *E*_*LHCGR*_=111.0%; figure 3), *GPER* and *RPS7* reactions were not (*E*_*GPER*_=71.1%; *E*_*RPS7*_=51.0%; figure 3). Moreover, the *r* value of *GPER* standard curve is low (r^2^=0.89), reflecting the poor amount of gene copies in the template and relatively high Cq values (supplementary table S1; supplementary table S2). Different results were found by ddPCR analysis, which identified good efficiency of all primers used, indicated by *E* values falling within the 100±20% interval (figure 3).

**Fig. 3.**
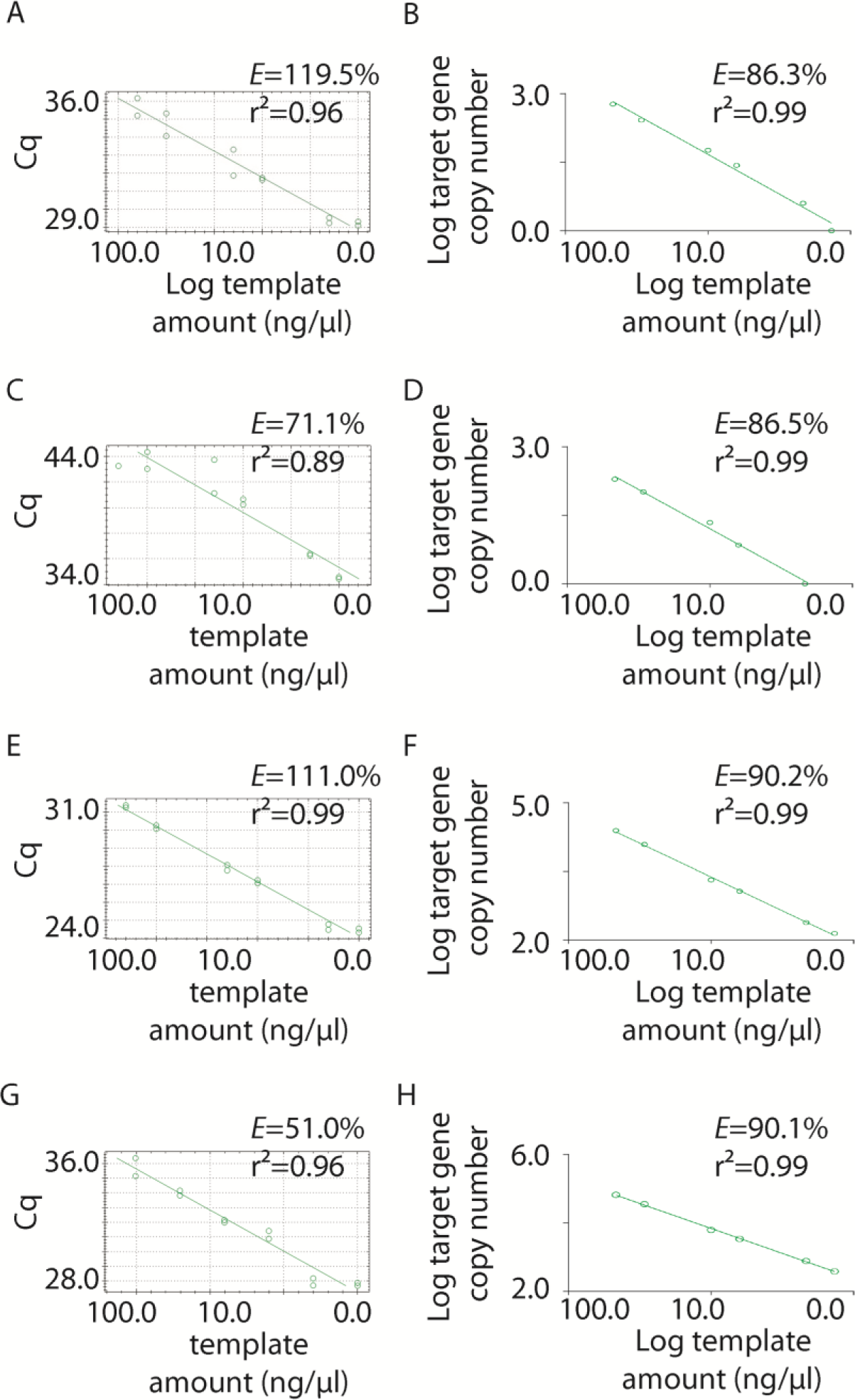
Evaluation of primer efficiency by qPCR and ddPCR. Serial dilutions of cDNA were prepared in nuclease-free water (100.0 - 0.5 ng/µl cDNA range) and plotted against qPCR and ddPCR amplification results (table 3) in a x-y graph. r2 and efficiency values were extracted from linear regressions used for interpolating data. A, B) FSHR efficiency data. C, D) GPER. E, F) LHCGR. G, H) RPS7. qPCR data are shown in the right panels, while ddPCR data in the left panels.

It is worth noting ddPCR analysis resulted in lower *LHCGR* expression levels than those of the *RPS7* gene, while opposite results were obtained by qPCR. These data might be due to 2.2-fold lower efficiency of *RPS7* than *LHCGR* primers using the qPCR method (figure 3).

### Calculation of transcripts number per cell

Finally, ddPCR results allow the calculation of transcript number per cell. Considering 17.50 pg/cell RNA (see methods section), as well as 100.00 ng/µl RNA (from 5714.29 cells) used for retrotranscription reactions with hypothetical 100% efficiency, 100.00 ng/µl of cDNA were obtained and used for ddPCR analysis in a total volume of 20 µl. Thus, the number of gene transcripts per cell is represented by the ratio between the “gene transcripts/µl” value (table 3), obtained using the cDNA concentration of 100.00 ng/µl, and 5714.29 as the relative cell number. Subsequent values of gene transcripts per cell were calculated after proportioning of the cell number with the corresponding cDNA concentration.

The following, approximate transcript numbers per cell were detected: *FSHR*=0.17±0.05; *GPER*=0.05±0.03; *LHCGR*=8.46±0.82; *RPS7*=24,52±2.05 (means±SD; n=6). Since these results were obtained as mean values from more than 5 thousand cells, transcript numbers <1.00 would be interpreted as a few positive cells admixed to negative cells.

## Discussion

Our results indicated that qPCR and ddPCR technologies are not similarly performing in detecting *FSHR, GPER, LHCGR* and *RPS7* gene expression, even if they both allowed this analysis. In hGLC, the observation of Cq values and transcript numbers revealed that *FSHR* and *GPER* were poorly expressed compared to *RPS7* (i.e. 0.7 and 0.2% of the housekeeping gene expression, respectively, detected by ddPCR), while relatively high *LHCGR* gene expression levels (34% of *RPS7* gene expression) were found by both qPCR and ddPCR (table 2; supplementary table S1; supplementary table S2). These genes are expressed at different levels in cultured hGLC (Myers et al., 2008), as well as in ovarian tissues (Heublein et al., 2014; Jeppesen et al., 2012), ovarian cancer cells (Heublein et al., 2013) and granulosa cell lines (Casarini et al., 2016a; Sasson et al., 2003). *LHCGR* transcripts per cell are about 50-fold higher than *FSHR* transcripts, which, in turn, are about 3.4-fold higher than the very poorly expressed *GPER* transcripts. These results are consistent with the luteinized state of the cells, as well as with the not clearly determined GPER functioning as a specific estradiol receptor that mediates endogenous estrogen effects *in vivo* (Luo and Liu, 2020). These data may lead to helpful insights in attempting to personalize the pharmacological treatment with exogenous hormones of patients undergoing assisted reproduction. The quantification of transcripts may be applied for studying pathophysiological conditions depending on hormone receptor activity, such as polycystic ovary syndrome or for predicting the normo- or poor-response of women to controlled ovarian stimulation, as well as in *in vitro* experiments for drug testing.

Our results indicated that, for equal amount of starting cDNA, ddPCR detects with high precision low copy number of gene transcripts, such as *GPER* and *FSHR* (table 3), even at sub-optimal annealing temperatures (table 2), while this is not the case for qPCR (table 3). These results were confirmed by relatively low correlation coefficients between transcript number and amount of template cDNA. Although these data are not surprising, since it is well-established that ddPCR has potentially higher sensitivity than qPCR (Hayden et al., 2013; Zhao et al., 2016), they suggest the use of ddPCR as a preferential method for gene expression screening of samples with low cell number or poor yield of DNA extraction. These data may be relevant for approaching personalized human infertility treatments, where the characterization of hormone receptor expression levels may be useful for predicting patient-specific response to the clinical hormonal treatment (Bosco et al., 2017; Simoni and Casarini, 2014). ddPCR and qPCR differ for the conditions in which primers work, resulting in different amplification efficiency of probe pairs. At the ideal efficiency of 100%, the amount of amplicons is doubling at each PCR cycle (Svec et al., 2015). The inconsistency of *FSHR* and *GPER* qPCR results, highlighted by their high variability (table 3, %CV), could be due to low primer binding affinity, likely exacerbated by low number of target transcripts in the template DNA and the potential presence of residual contaminants remaining after nucleic acid extraction procedure and subsequent manipulations. These limitations are surpassed by ddPCR technology, which is based on the detection of fluorescent signals, emitted after end-point PCR amplification of each single transcript copy and partitioned in single droplets (Beer et al., 2008). This method allows the determination of the number of transcripts through the count of positive/negative events, determined by the presence or absence of light emission after properly establishing the signal amplitude intensity threshold. The better performance of ddPCR compared to qPCR allows the detection of signals due to unspecific primer targeting in the *RPS7* housekeeping gene. The presence of these transcripts could be due to excessive cDNA template, likely increasing the presence of potential contaminants, which leads to unspecific primer binding. These signals are detected as a sub-population of positive droplets falling above the fluorescence amplitude intensity threshold, but having lower signal amplitude intensity than those where the target is correctly determined. The *RPS7* gene, encoding the 40S ribosomal protein S7, is usually used as a reference for gene expression analysis in several research studies (Armakolas et al., 2012; Kessler et al., 2009; Kim et al., 2019), including those falling in the context of human fertility (Casarini et al., 2017, 2016a, 2016b). Therefore, these findings are relevant for optimizing the amount of cDNA template in gene expression analyses and avoid biased results due to wrong data normalization.

In conclusion, these two methods are based on different approaches and lead to overall similar results. However, ddPCR allows a clear determination of unspecific binding and sub-optimal performance of primer pairs, generally due to low target expression, improper amount of template cDNA or contaminants. Indeed, the absolute number of *LHCGR* transcripts per cell were determined by ddPCR, but not by qPCR, revealing they are about 50-fold higher than *FSHR* and 170-fold higher than *GPER* transcripts per cell. These conditions must be carefully assessed before hormone receptor gene expression analysis by qPCR, in order to achieve an accurate evaluation of hormone receptor’s transcriptional profile in patients undergoing infertility treatment or affected by reproductive disorders.

## Acknowledgements

This study was performed in the context of the Departments of Excellence Programme supported by MIUR and granted to the Department of Biomedical, Metabolic and Neural Sciences of the University of Modena and Reggio Emilia.

## Authors contact information

Samantha Sperduti, Phone: +39.0593961713, Fax: +39.0593962018;

Claudia Anzivino, Phone: +39.0593961705, Fax: +39.0593962018;

Maria Teresa Villani, Phone: +39.0522296466, Fax: 0522335200;

Gaetano De Feo, Phone: +39.0522296466, Fax: 0522335200;

Manuela Simoni, Phone: +39.059-3961815, Fax: +39.0593961335;

Livio Casarini, Phone: +39.0593961705. Fax: +39.0593962018;

