## supplementary material for "Comparison of real-time quantitative Polymerase Chain Reaction (PCR) and digital droplet PCR for quantification of hormone membrane receptor *FSHR, GPER* and *LHCGR* transcripts in human primary granulosa lutein cells"

**Supplementary Fig. 1** Annealing temperature-dependence of target sequence amplifications by qPCR and ddPCR. Pictures are output images of *FSHR* (A, B), *GPER* (C, D), *LHCGR* (E, F) and *RPS7* (G, H) melting and gene expression analysis obtained by qPCR (left-panels) and ddPCR (right-panels) using a CFX96™ Real-Time PCR Detection System and a QX200 Droplet Digital PCR (ddPCR™) System (Bio-Rad Laboratories Inc.), respectively. Reactions were prepared from the same PCR mix divided for the analysis in duplicate using the following annealing temperatures: 56, 58 and 60°C. NTC=negative control samples.
